# Targeting phosphorus loss with carbon farming practices? Results from an on-farm study

**DOI:** 10.1101/2025.09.01.673431

**Authors:** Tuomas J. Mattila, Jari Niemi

**Affiliations:** Finnish Environment Institute SYKE, Climate solutions unit

## Abstract

Agricultural soils lose P to waterbodies and C to the atmosphere. Reversing the C trend requires change in management (carbon farming), but what is the effect of carbon farming practices on P transport from soils to waterbodies? This was empirically studied by analyzing the P loss risks from 20 farms participating in a 5-year on-farm carbon farming experiment. We evaluated the effect of C farming practices on soil P balance, P stocks, and potential P loss through surface and subsurface runoff and erosion. Based on the results, carbon farming had only minor impacts on P loss compared with the current soil conservation practices already applied by the farms. The farmers did not utilize the opportunity to reduce P fertilization through some carbon farming practices which could improve soil P availability, therefore resulting in weak P balances. Furthermore, only a small fraction of the field area (18%) was responsible for the majority (50%) of the estimated P loss, indicating the importance of P loss hotspots. C farming practices do not seem to improve water quality unless they are tailored to target key processes of P loss such as maintaining a negative P balance on high-P sites, reducing runoff, and focusing on local critical source areas.

## 1. INTRODUCTION

Agricultural soils are the main source of phosphorus (P) loading to water bodies (Fink et al., 2018) and a major source of greenhouse gas emissions (Li et al., 2025). The climate impact can be mitigated by changing farming practices to sequester more C (Paustian et al., 2019; Smith et al., 2019). The main C sequestration methods rely on two key processes: i) increasing the vegetative period to capture more C and ii) avoiding soil disturbance to reduce C loss (Chenu et al., 2019; Mattila and Vihanto, 2024; Paustian et al., 2019). As these processes are also widely used for soil conservation, could carbon farming practices help solve water quality issues from P loss?

Many synergies exist between C sequestration and P loss mitigation (Table 1). Carbon farming practices, such as cover crops, grass in crop rotation, or improved grazing, also reduce erosion (Duncan et al., 2019; Franzluebbers et al., 2012; Kleinman et al., 2022), which is the dominant P loss pathway (Kronvang et al., 2007). Practices that increase the vegetative period also increase evapotranspiration thereby reducing P loss via runoff (Hanrahan et al., 2021). Maintaining soil residue cover improves infiltration and reduces runoff and erosion (Panagos et al., 2015). In theory, multiple benefits could be obtained from applying C sequestration practices.

**Table 1.**
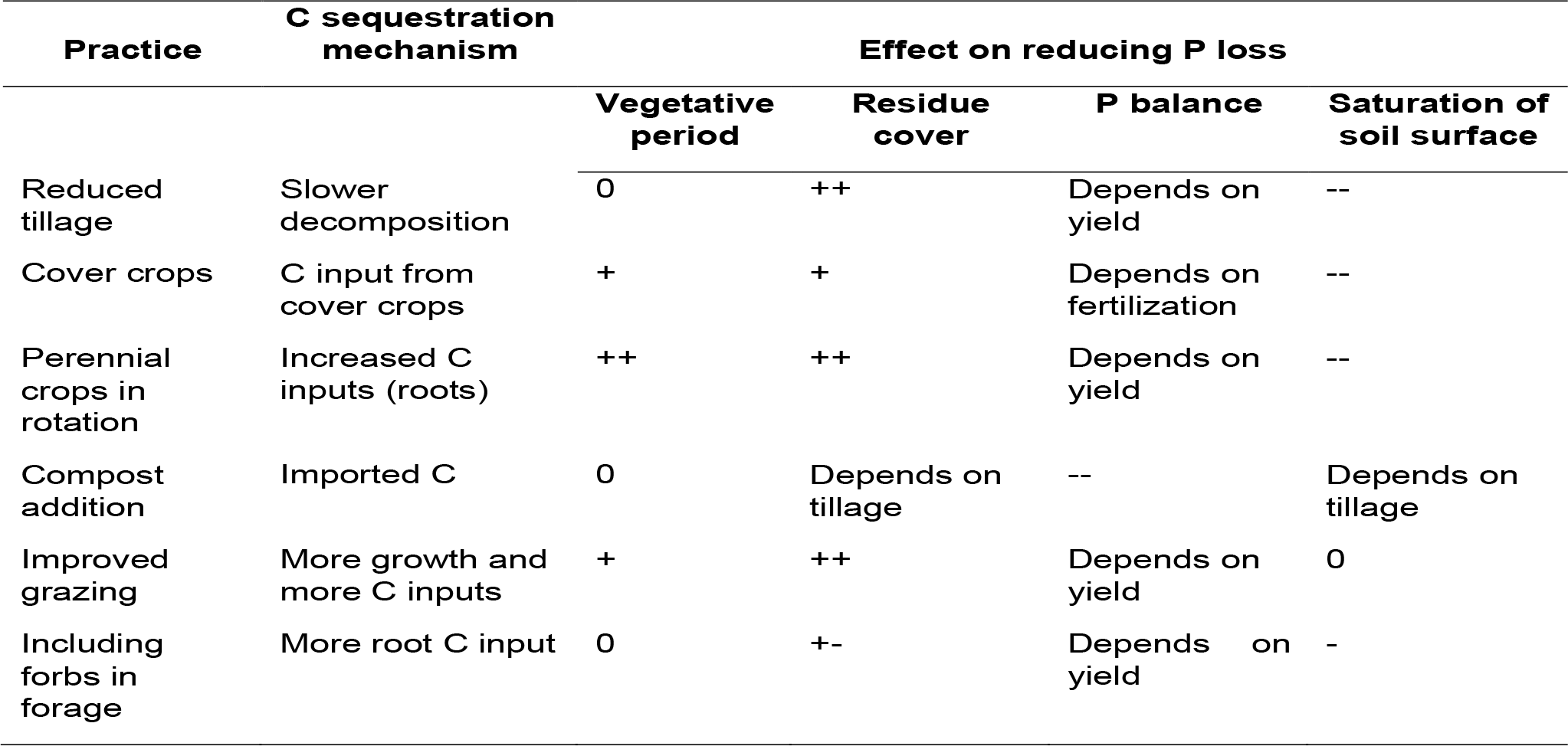
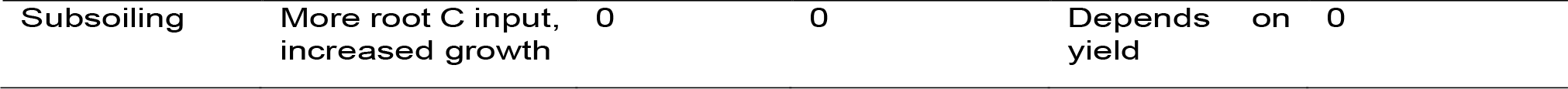
The potential of carbon farming practices to mitigate P loss. In this study, reduced tillage was included in the business-as-usual control, and the other measures were considered as additional carbon farming practices.

However, in practice soil conservation practices can increase the loss of highly bioavailable dissolved P (Duncan et al., 2019; Jarvie et al., 2017; Uusitalo et al., 2018). Agricultural soils have accumulated vast reserves of legacy P as a result of fertilization practices in the previous century (Mattila, 2024; Shober et al., 2025). When the accumulated P exceeds a critical fraction of the soil’s sorption capacity (approximately 10% of Fe and Al), P becomes more available, resulting in increased loss of dissolved reactive P (DRP) (Kleinman, 2017; Nair et al., 2020). When the soil is not mixed by tillage and P-containing plant residues accumulate on the soil surface, the sorption capacity in the soil surface is often exceeded, increasing DRP loss (Jarvie et al., 2017). Reducing DRP loss requires a negative P balance (fertilizer in vs. harvest out) to “mine out” accumulated P to a level below the DRP threshold, which can take decades (Nair et al., 2020; Wall et al., 2013). Carbon farming practices can either help or harm achieving negative P balances. Reduced tillage can also reduce yields (Pittelkow et al., 2015; Uusitalo et al., 2018), thereby slowing P removal. Cover crops can improve P reserve availability for later crop use (Hallama et al., 2019). However, if the fertilizer rates are not adjusted subsequently, this solubilized P becomes a risk for DRP runoff (Cober et al., 2019; Maltais-Landry and Frossard, 2015). The extent to which the P solubilized by cover crops becomes DRP loss also depends on the local soil conditions (Cober et al., 2019), highlighting the need to customize P management on a field-by-field basis and to focus on high-risk fields (Mattila, 2024).

Evaluating the P loss from a field is challenging because most P loss occurs during brief peak flow events (Hanrahan et al., 2021). Capturing the peak requires repeated sampling of both surface runoff and subsurface drainage, which is feasible mainly for instrumented experimental fields (Uusitalo et al., 2018). The other option is to measure riverine P loading from the whole catchment, tracking management effects at the landscape level (Ekholm et al., 2024; Hanrahan et al., 2021; Shober et al., 2025). Unfortunately, catchment-level analysis cannot locate the source of the P loss comes from, especially if there are P loss hotspots (critical source areas, CSAs), which occupy a minor area but produce most of the P loading (McDowell et al., 2025). The P index is the most widely used tool for identifying CSAs. It combines P source (soil P, fertilization practices) and P transport (erosion, waterflow patterns, connectivity to waterbody) into a relative risk score (Osmond et al., 2025). Most current P indices are split to separate flows for erosion, surface runoff, and drainage (Osmond et al., 2017, 2025). The P indices correlate well with actual P loss (Osmond et al., 2017; Thomas et al., 2016) and match modeled P loss (Bolster and Vadas, 2022; Fiorellino et al., 2017). Overall, P indices can identify the relatively few fields (5-20%) that contribute to the majority of P loading and guide site-specific P loss mitigation practices (McDowell et al., 2025).

In this study, we analyzed the tradeoffs and synergies of C sequestration and P loss across different farms. We used an on-farm experiment (Mattila et al., 2022), where the farmers co-designed and maintained a split-field experiment testing various C sequestration practices (Table 1). Comparing carbon farming with business-as-usual enabled the analysis of the effect of adding carbon farming to existing farming practices. We estimated P loss at the start of the experiment and after five years of different practices by integrating high-resolution erosion mapping, on-site measurements, soil samples, and farmer notes. We then evaluated the extent to which P loss occurs from specific hotspots and the influence of carbon-farming practices on P loss through erosion, P balances, and runoff. Finally, we evaluated the potential for improving carbon farming practices to achieve simultaneous improvements in C sequestration and P loss.

## MATERIALS AND METHODS

### 2.1 Study sites and data sources

We studied a subset of 20 intensively monitored farms chosen from the 105 participants of the Carbon Action carbon-farming experiment (Mattila et al., 2022). The subset of farms was chosen to cover different farming systems, soil textures, and geographical locations. The farms were distributed in an area of 200 × 500 km covering the main agricultural areas of Finland (ranging from 63.18 N to 60.36 N in latitude). As such, the farms were a representative sample of the early-adopter farms and could be used to assess how the increased use of carbon farming practices influences P loss across Finland.

Each farm had an experimental field (2-5 ha), where carbon farming was applied on one half of the field and the other was kept as a business-as-usual control. Both field-half plots had three monitoring points fitted with continuous soil moisture and temperature sensors (Meter Group Teros-12 capacitive sensor). Monitoring points were visited each year in July for soil infiltration measurement, structure assessment, and soil sampling. Soil samples (30 samples per field half, 17 cm depth) were used to measure soil P, Fe, and Al using Mehlich-3 extraction (Mehlich, 1984). These concentrations were used to estimate the P sorption capacity and degree of P saturation (DPS; molar ratio P / (Fe + Al) (Dari et al., 2018; Kleinman, 2017)). Additional soil samples were also used to measure bulk density, soil texture, and soil organic matter (SOM), which were then used to estimate soil porosity and water holding capacity and also as input in the APLE phosphorus model. Details of the sampling and sample processing are given in the published data repository (Mattila et al., 2023).

The on-site monitoring was complemented with publicly available rainfall (Finnish Meteorological Institute, 2025) and soil erosion susceptibility (Natural Resources Institute Finland, 2023; Räsänen et al., 2023) datasets. The high resolution (2×2 m) erosion susceptibility map was based on the RUSLE soil loss equation and its factors related to rainfall (r), slope (l and s), and soil type (k), but cropping (c) and erosion mitigation measures (p) were not included. We added them based on regional factors (Räsänen, 2024) and cropping and tillage practices documented in the farmer notes. The farmer notes also included fertilization amounts, methods, and yield, which were used to calculate P balances for the plots (Panagos et al., 2022).

### 2.2 Estimating the P loss

#### 2.2.1 Dissolved P loss risk

We used the DPS to identify fields with excessive P levels and high dissolved P loss risk (Dari et al., 2018; Kleinman, 2017; Nair et al., 2020). In brief, the molar soil phosphorus concentration was compared with the sorption capacity for P to calculate a degree of phosphorus saturation (DPS = P/Fe + Al). If DPS was > 10% (Dari et al., 2018), the field was considered to have a high risk of dissolved P loss.

#### 2.2.2 P loss through runoff and erosion

A simplified water balance was applied to evaluate the short-term effects of carbon farming practices on runoff (Mattila, 2024). Rainfall was divided into runoff (surface and subsurface) and temporary soil storage. Soil storage occurred when the soil was below field capacity (volumetric water content smaller than water holding capacity, VWC < WHC) and the rain intensity was lower than the soil surface hydraulic conductivity. We measured hydraulic conductivity with a mini-disk tension infiltrometer (Meter Group, USA; 15 mm suction, 2 mm sand layer to improve soil contact) with five replicates on each plot. The infiltration rate was converted to hydraulic conductivity by fitting cumulative infiltration vs. time and using van Genuchten parameters specific to soil texture (Dohnal et al., 2010; Zhang, 1997). Hydraulic conductivity was used to identify rainfall events which would result in surface runoff due to high rain intensity.

Runoff or drainage would also occur with low-intensity rain if the soil was already saturated. Rainfall on non-saturated soils could be quantified as effective rainfall (Brouwer and Heibloem, 1992). First, field capacity (FC) was estimated using pedotransfer functions based on soil texture and SOM (Szabó et al., 2021). FC was then adjusted to match the observed water content in each field in early spring. This resulted in a slight (average 7%) adjustment for 19 of the 40 plots, reflecting the effect of soil structure on water holding capacity (Koudahe et al., 2022). The time series was filtered for periods when the water content was less than the FC and peak events corresponding to rainfall were identified (Bean et al., 2018). Peaks were defined as local maxima within a 72-hour period and the effective rainfall as the increased water storage compared to 48 hours before the peak. To avoid counting water that would run off or drain, the peak maximum was reduced to FC before calculation.

The RUSLE erosion susceptibility map was based on soil type-specific erodibility factors (k) (Räsänen et al., 2023). The soil type maps may misclassify individual soils, and the soil type classes cover large ranges of clay content (20-80% for Stagnosols), which results in large errors for soils which at the extreme ends of the range. We used measured soil texture and infiltration to improve the erodibility assessment, using a recent methodological improvement (Gupta et al., 2024). In addition to a more accurate assessment of erodibility, this enabled the evaluation of how erodibility developed with changes in hydraulic conductivity over time.

Local factors were used to include the RUSLE reduction effects of crop (annual 0.4, perennial grass 0.3) and tillage (1 inversion tillage, 0.76 minimum tillage, 0.44 no-till, 0.36 perennial crop) (Räsänen, 2024). In addition, cover crops were credited with a 20% erosion reduction (Panagos et al., 2015). Finally, when the field had a functioning drainage system, erosion was reduced by 40% (P factor) (Räsänen, 2024), but unfortunately, many of the fields had drainage problems (Mattila and Vihanto, 2024). In effect, cropping practices could have a considerable effect on erosion; given a certain erosion susceptibility, the erosion could range from 0.4 (annual crop with inversion tillage) to 0.11 (perennial crop over winter).

#### 2.2.3 P-indices and APLE

The results from erosion, site measurements, and fertilization practices were integrated into two P indices and one P loss model (APLE). The first P index was the classical source x transport-index based on Sharpley (Osmond et al., 2025; Sharpley et al., 1993) but updated score values were used (Weil and Brady, 2016). The second P index was based on the recommended development of splitting the source and transport factors into separate flows (i.e. erosion, surface runoff, and drainage) (Osmond et al., 2017, 2025). We used the most recently updated P index of this type, the Iowa P index (NRCS, 2024), which estimated the erosion component from erosion (RUSLE), sediment delivery (distance to waterbody and type of landform), possible buffer zones, residue management (stratification) and soil test P (Mehlich-3). The runoff component was estimated from the precipitation, runoff factor, soil test P, and P fertilization rate and method. Finally, the subsurface drainage component was estimated from precipitation, drainage factor, and soil test P, and then the three flows were combined to obtain one P loss index (NRCS, 2024).

The APLE 3.0 (Bolster and Vadas, 2022) also simulates P loss from different flows but excludes subsurface drainage and splits the runoff into dissolved and particulate P loss. The particulate P loss is based on erosion and P enrichment into the eroding layer. The dissolved P loss is based on the fertilizer and manure application rates as well as the dissolved P loss from the accumulated soil P. Precipitation, runoff properties (curve number), tillage (mixing), fertilizer placement and soil properties (texture and OM) influence these processes providing a good basis for site-specific estimation of P loss.

### 2.6 Statistical analysis

The statistical analysis focused on whether there was a change over time in the plots between 2019 and 2023 (Δ_N_=N_t_ – N_t-1_) and especially if the change would be different between the carbon farming and business-as-usual (Δ_Ntreatment_ - Δ_Ncontrol_). As the experiment was a pair-wise comparison, with each farm as a separate experimental block with randomly assigned treatments, a paired analysis was used for statistical testing. Since P loss is a result of multiplicative processes, a normal distribution cannot be assumed, a non-parametric Wilcoxon signed rank test was used to evaluate the significance of the difference. We used linear models (least squares regression) to test the magnitude of the effects and the significance of correlations with explanatory variables. The coefficient of determination (R^2^) was used to evaluate the strength of the correlations. All statistical analyses were conducted using the R software (R Core Team, 2022).

## 3. RESULTS

### 3.1 P loss risks at the start of the experiment

The erosion was low but highly variable (Figure 1): the mean soil loss was 0.71 t/ha/yr, but it ranged from 0.03 to 7.78 t/ha/yr between plots. The erosion susceptibility (r x s x l) term determined 85% (R^2^=0.85) of the variability between fields, the remaining 15% was controlled by soil erodibility (k). Crop coefficients (c) and erosion mitigation practices (p) had only a minor effect in this dataset, as crop coefficients were low across all fields (mean 0.16 ± 0.05) due to the widespread use of conservation tillage practices.

**Figure 1.**
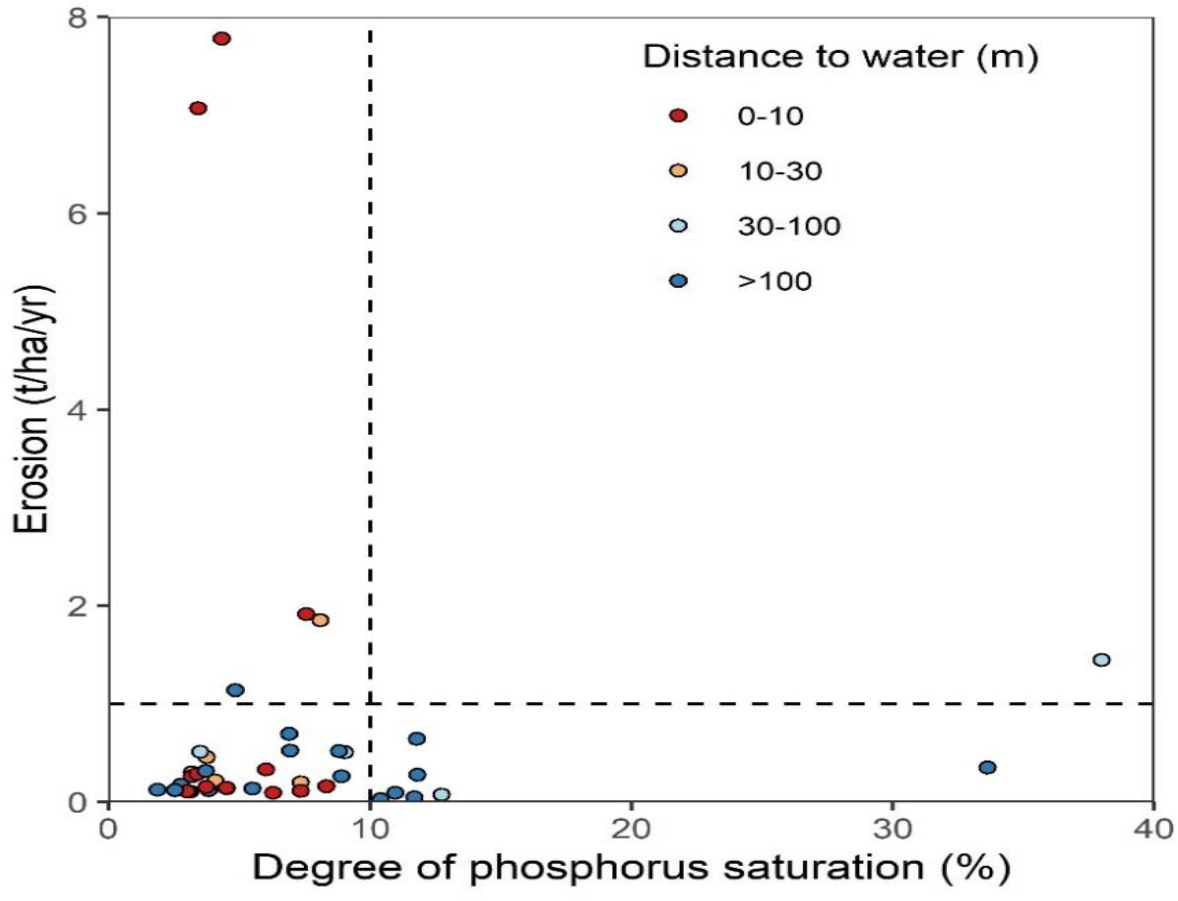
Phosphorus loss risks through erosion and excess P saturation in the studied fields at the start of the experiment. The dashed lines are thresholds for erosion (soil formation rate) and degree of P saturation (10%).

**Figure 2.**
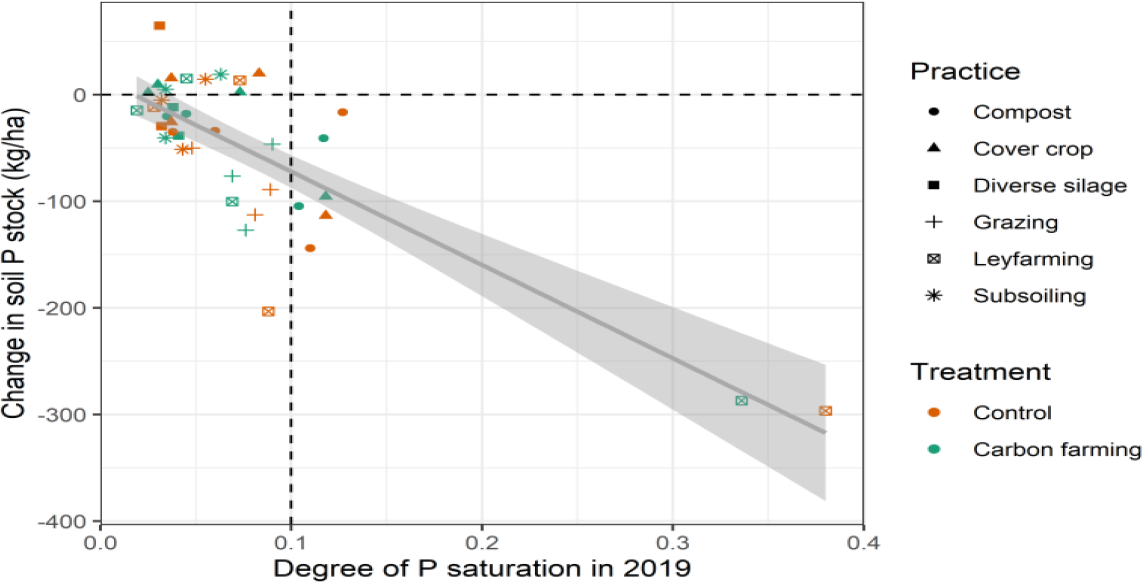
Change in soil P stock (Mehlich-3 extracted P) in relation to starting P status. The dotted vertical line describes a risk level for P saturation.

The erosion was not evenly distributed within the plots. Comparing the erosion values with the thresholds for sustainable, critical and severe erosion (Panagos et al., 2015): 13% of the area had an erosion rate faster than soil formation (1 t/ha/yr), 4% exceeded the 5 t/ha/yr critical limit, and only 1.6% exceeded the limit for severe erosion (10 t/ha/yr). Almost all (99%) of the severe erosion was found on one field, which had a steep long slope and therefore had severe erosion on 24% of the area. In contrast, for most plots (65%) only 1% of the area had erosion rates higher than the soil formation rate indicating that erosion problems were localized within and in specific areas of individual fields.

The phosphorus saturation (DPS) was also unevenly distributed (Figure 1). Eight of the 40 plots (20%) exceeded the DPS threshold of 10%. The average DPS was 7% (± 7%). The average sorption capacity (Fe+Al) was 45 mmol/kg (±10 mmol/kg) and ranged between 28-68 mmol/kg. However, P concentration (30-430 mg/kg) controlled most of the DPS variability (R^2^=0.93). The DPS was exceeded when the P concentration was > 140 mg/kg. There was also a strong correlation between DPS and the Finnish acid ammonium-acetate extraction (R^2^=0.95), where a P concentration of > 18 mg/dm^3^ corresponded to a DPS of > 10%. The Finnish AAc result showed weaker correlation with Mehlich-3 P (R^2^=0.85); therefore, it could be thought to represent a highly available P pool similar to that of the DPS. Regardless of whether DPS or the Finnish labile P measurement was used, a small fraction (15-20%) of the fields was identified as a potential risk for dissolved P loss.

The fields also differed in their P transport properties. The average distance to the water bodies was 140 m, but it ranged between 2 and 550 m. The worst combination was in plots with high erosion and proximity to water (10% of plots). The multiplicative P index and the component-flow P index weighted these risk factors differently resulting in different prioritizations. The multiplicative index did not prioritize plots at all: all fields were in the medium risk category (50-250 points). The variation in multiplicative P indexes was controlled slightly more by the transport factor (R^2^=0.41) than by the source factor (R^2^=0.33). The highest risk scores were on plots with high connectivity to waterbodies (close distance, no buffer between field and drainage ditch) and high manure use. The component-flow index effectively prioritized the plots: 15% of the plots had medium risk and 2.5% had high risk. The highest P indexes were on plots with high P saturation, and the next highest indexes were on plots with the highest erosion rates. On average, surface runoff was the dominant loss pathway (72%), followed by erosion (17%) and subsurface runoff (11%); however notable differences were observed, especially when the erosion rate was high. As the factors were correlated through shared factors, such as P concentration, surface flow explained 56% of variability (R^2^=0.56), erosion 51% and subsurface runoff 21%.

The APLE 3.0 model predicted an average particulate P loss rate of 1.51 kg/ha/yr (±1.55 kg/ha/yr). The highest particulate P loss (6.93 kg/ha/yr) was estimated for the previously mentioned high-erosion plot. The average DRP loss was much lower (0.03 ± 0.03 kg/ha/yr), and the highest DRP loss (0.13 kg/ha/yr) was from a plot with low permeability (high runoff) and very high P concentration in the soil. The plots with the highest P loss had approximately four times the average P loss. Expressed as a Pareto curve: 18% of plots amounted to 50%, and 45% amounted to 80% of the estimated total P loss. When considering only the DRP, the same relationships held (18% of the plots amounted to 50% of load; 38% amounted to 80% of the load), but the priority plots were different. The dissolved P losses were highest in the perennial silage grass fields where fertilizer was surface-applied, and in the fields with a high starting P concentration and low permeability.

### 3.2 Farming practices related to P loss

Farmers applied six carbon farming practices (cover crops, grass in rotation, improved grazing, forages in grass, subsoiling, and soil amendments), which influenced P loss through changes in soil cover, tillage, fertilization, P balance, and hydrology.

The carbon farming practices had only a minor effect on soil tillage and subsequently soil cover, as the farmers already applied conservation tillage practices. On the control plots, of the 100 site-years in the study, only 1% had inversion tillage with no soil cover, and 4% had mulch tillage. More than half (52%) had vegetative cover over winter and the remaining (43%) had undisturbed crop stubble. Although some farms (40%) applied inversion tillage (mouldboard plowing) in rotation, they either applied it in spring prior to crop seeding or in late summer before establishing a winter crop, thereby maintaining soil cover during winter season, when the soil was erosion prone. Two farms (10%) applied no-till, and three farms (15%) had perennial grass. Carbon farming practices mainly replaced crop stubble with living plants, resulting in 86% of the soil cover being vegetative cover and 10% crop stubble in the carbon farming plots. In addition, grass in rotation reduced tillage by replacing an annual crop with a perennial crop.

In theory, cover crops and grass in rotation could reduce P fertilization by improving P availability to the subsequent crops. However, the farmers did not adjust fertilization based on cover crops, but the cereal farms with grass in rotation left the grass crop unfertilized, thereby decreasing the total amount of P applied. The farmers also did not adjust their fertilization practices based on the soil P level (R^2^=0.07, p=0.12). Overall, the difference in fertilization between carbon farming and control was minor and non-significant (9.7 vs. 10.2 kg P/ha/year, p=0.26), but including grass in rotation decreased P application by −4.2 kg/ha/yr (p=0.09) and compost application increased it by +2.6 kg P/ha/yr (p=0.06). Farmers applying compost reduced the mineral P fertilizer application, resulting in more P being sourced from organic sources in the carbon farming plots (49% vs. 42%). As most farms left the grass in rotation unharvested, the effect on P balance was weak. Overall, the carbon farmed plots had a less-negative P balance than the control group (−1.2 vs. −2.0 kg P/ha/yr, p=0.01) indicating slower P removal. For compost application, the difference between carbon farming and control (+1.6 kg P/ha/yr) was due to increased P application, for grass in rotation (+0.7 kg P/ha/yr) due to reduced P removal with crop, and for cover crops (+0.5 kg P/ha/yr) due to lower crop yields in one of the experimental farms. Overall, 30% of the plots had a positive P balance, associated with the application of manure. The cereal cropping farms without repeated manure application had a negative P balance ranging from −2 kg P/ha/yr to −11 kg P/ha/yr. The largest negative P balance (−16 kg P/ha/yr) was observed on a farm, that did not apply manure but harvested the rotational grass crop.

In principle, increasing vegetative cover could also influence the water balance of soil, by keeping the soil drier and allowing more temporary storage of water. This was analyzed from the time series of on-site soil moisture sensors and rainfall data. On average, the plots were wetter than field capacity for 35% of the time, and this fraction was similar for both carbon farming (34%) and control plots (36%). However, some carbon farming practices had significant differences in the duration of wet soil conditions compared to control: grazing (+10%, p=0.03), leyfarming (+5%, p=0.06), and subsoiling (+10%) tended to increase the time periods with wet soil, while compost application decreased it (−12%, p=0.06). Cover crops had a minor (−4%), but variable (p=0.65) effect. Although the soil was wetter than field capacity for a third of the time, approximately half of the rainfall (50% ± 13%) occurred during this time (autumn-winter), resulting in considerable runoff. Compost and cover crops decreased runoff, whereas grazing, leyfarming and subsoiling increased it, however, these changes were not systematic or statistically significant, possibly due to high variability in soil conditions and slope position between control and carbon farming in individual sites (i.e., the randomly assigned treatment was either upslope or downslope, increasing variability). Based on the observed peaks in soil moisture during rain events in the growing season, on average 15% ± 7% of annual rainfall (105 mm out of 700 mm) was temporarily stored into the topsoil for later plant use (effective rainfall). Again, no systematic, statistically significant difference was observed between carbon farmed and control plots, but compost and cover crops slightly increased effective rainfall, while grazing, leyfarming, and subsoiling decreased it.

### 3.3 Effect on P loss after 5 years

Carbon farming practices may reduce P loss through i) reduced soil P availability (dissolved and particulate P) or ii) reduced erosion (particulate P). We analyzed these processes separately and then combined the assessment using P indices and APLE model estimates.

The P concentrations in the soil decreased on average in both the carbon farming and control plots and they decreased the more, the higher the starting P concentration was (p < 0.001, average relative decrease 12% in 5 years). Soils with low starting P concentrations gained P; however, when the levels were agronomically excessive (>100 mg/kg P), the P concentrations decreased in all the observed plots. The decrease was insufficient to transition fields with high starting phosphorus saturation (DPS >10%) to a low-risk state. In addition, one pasture plot and one plot with chicken manure applications exceeded the 10% threshold during the experiment.

Carbon farming did not significantly influence the change in phosphorus concentration or saturation. The change over time also did not correlate with the P balance (fertilization – yield removal), but it correlated strongly with the starting P saturation (R^2^=0.65, p<0.001). In general, soils with higher starting P levels lost more P than soils with low starting P levels. Generally, a 5% increase in starting DPS resulted in an average annual decrease of 9 kg/ha/yr. This relationship was maintained even when one farm with extreme DPS (>30%) was removed from the dataset. This highlighted the influence of soil P levels in contrast to that of farming practices or P balances when considering future changes in soil P concentrations.

Erosion decreased between 2019 and 2023 in both the carbon farming and control plots (0.06 t/ha/yr and 0.10 t/ha/yr; p < 0.01) (Fig 3a) as a combination of improved infiltration (14 mm/hr; p < 0.0001) and increased vegetation cover. The erosion was smaller in the carbon farmed plots than in the control both at the start of the experiment (−0.15 t/ha/yr, p=0.02) and at the end (−0.07 t/ha/yr, p=0.03). The erosion did not decrease more in the carbon farming plot than in the comparative control plots, indicating that the erosion reduction benefit was achieved already at the beginning of the experiment and did not improve with the duration of the experiment.

**Figure 3.**
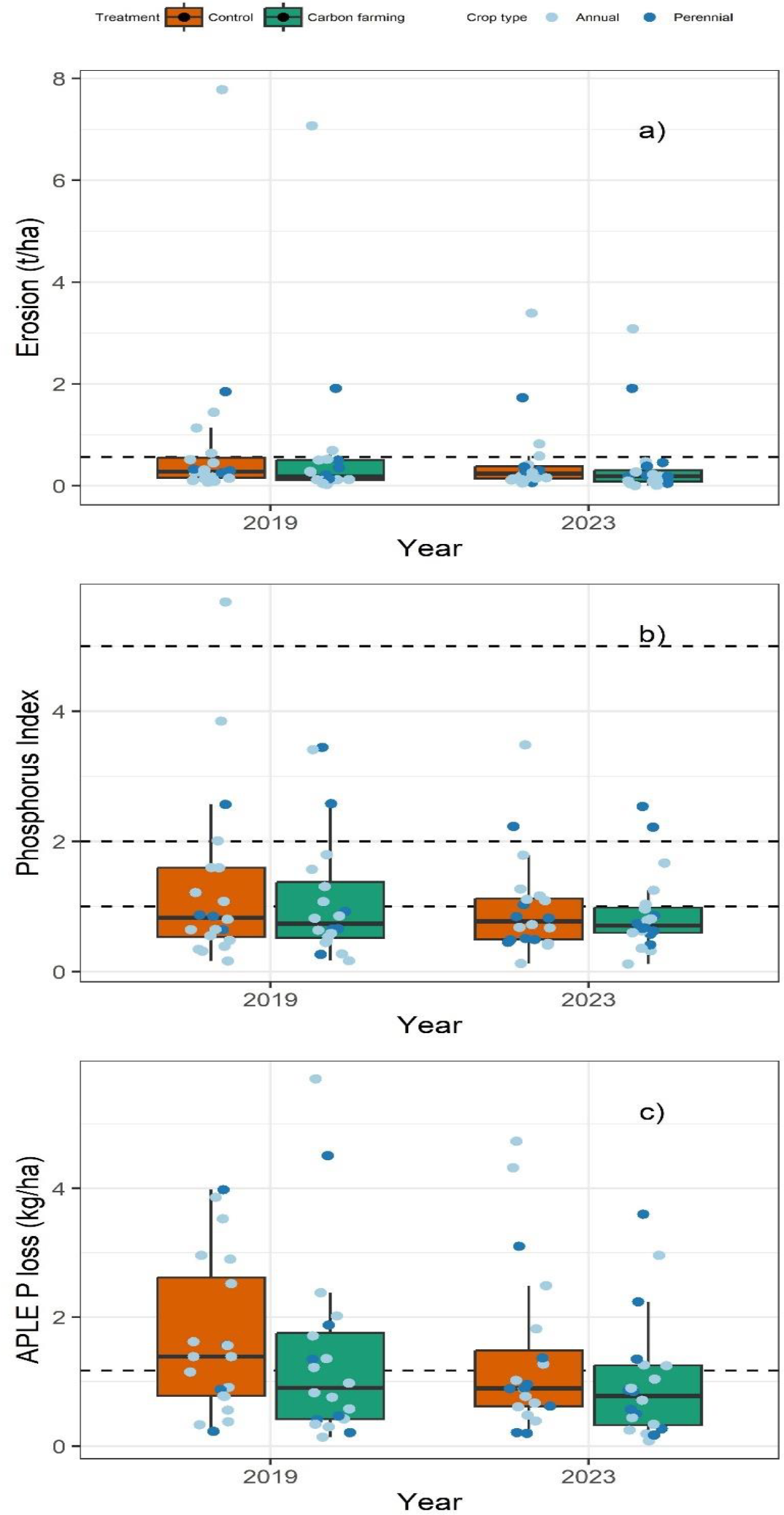
Changes in erosion, phosphorus index and estimated P loss between 2019 and 2023 in carbon farming and control plots.

Reduced erosion also resulted in a smaller component-flow P index in 2023 than in 2019 (p=0.006, average 0.28 index points) (Fig 3b). However, the P index decreased more in the control plots than in the carbon farming plots (0.09 index point difference, p=0.05) resulting in a smaller P index in carbon farming at the start of the experiment (p=0.05), but no statistical difference at the end of the experiment between the treatments.

The APLE P loss (Fig 3c) systematically decreased in the carbon farming plots (p=0.01, average 0.38 kg/ha/yr), but not in the control plots (p=0.06). This resulted in smaller P loss from carbon farming plots than from the corresponding control plots at the end of the experiment (p=0.04, −0.39 kg/ha/yr), especially for cover crops (−0.6 kg/ha/yr) and leyfarming (−1.1 kg/ha/yr), which increased vegetative cover. However, as the estimated P loss decreased on both carbon farming and control plots, the additional decrease from carbon farming practices was not statistically significant when compared across all practices.

## 4 DISCUSSION

Based on the results, the carbon farming practices had very little impact on P loss. This was especially true when they were added to current farming practices, where i) soil conservation measures were the norm and ii) the P balance was already negative. This was emphasized by on-farm study participant selection, as they were early-adopter farmers with strong interest in soil conservation.

Carbon farming and soil health measures such as cover crops have been found to reduce P loss through reduced erosion and runoff (Duncan et al., 2019; Hanrahan et al., 2021). In this study, the farmers maintained residue cover over winter also in the business-as-usual control plots, and carbon farming practices only replaced residue cover with vegetative cover. Although this resulted in small decreases in erosion, it was insufficient to cause a systematic difference between the carbon-farmed and control plots. In theory, increased vegetation cover should increase evapotranspiration and reduce runoff (Hanrahan et al., 2021), but we did not find evidence of this when analyzing the time series of rainfall vs. infiltration or soil moisture. In contrast, in some cases, including cover crops and grass in rotation increased runoff and decreased porosity due to reduced tillage and subsequent soil settling.

The risks of P loss decreased in both carbon-farmed and control plots due to gradual decreases in high P concentrations. Agricultural soils have accumulated considerable amounts of legacy P (Mattila, 2024; Panagos et al., 2022), resulting in a high risk of dissolved P loss (Kleinman, 2017; Nair et al., 2020). Removing excess P requires maintaining a negative P balance over several years to decades (Wall et al., 2013). In theory, increased vegetative cover could solubilize the legacy P to plant-available forms (Hallama et al., 2019; Maltais-Landry and Frossard, 2015), thereby reducing the need for P fertilization and helping to maintain the strongly negative P balances needed for rapid P removal. In practice, the farmers in this experiment did not adjust P fertilization based on cover crops. The rotated grass crops (leyfarming) were usually not fertilized, but the subsequent crop received the same amount of P regardless of previous year’s crop history. Farmers also did not adjust the fertilizer amounts based on the soil P concentration. Adjusting P fertilization to the availability of P to crops influenced by soil P status (Nair et al., 2020; Recena et al., 2022) and P solubilization from previous cover crops (Hallama et al., 2019) presents a key opportunity for accelerating the removal of accumulated legacy P.

As this carbon farming experiment was not conducted to mitigate P loss, several key processes and opportunities were left unused. In addition to reducing fertilizer amounts to match crop needs, a stronger focus on soil quality and within-field hotspots would have been useful. Based on a soil quality survey in the beginning of the experiment (Mattila and Vihanto, 2024), most of the fields were compacted and many had malfunctioning drainage systems. These factors both increase surface runoff and erosion as well as decrease yields and, therefore, P removal. A malfunctioning drainage system concentrates water to parts of fields, creating hotspots of local runoff and P loss. Overall, our results supported earlier findings, where a minority of the fields caused the majority of P loss (McDowell et al., 2025; Thomas et al., 2016). The actual fractions in different catchments vary between 2-50% of fields causing 20-90% of P loss (McDowell et al., 2025). In our data, 18% of area caused 50% of P loss and 40% caused 80% of P loss, which fits the previously published range. This supports the considerable potential to increase P loss mitigation efficiency by targeting sites with high erosion risk, high P concentrations, and strong connectivity to waterbodies (Mattila, 2024; McDowell et al., 2025; Thomas et al., 2016).

To better target hotspots, land managers should be able to first identify them. P indices have a long history of prioritizing sites for P loss (McDowell et al., 2025; Osmond et al., 2025). In our results, the classical multiplicative index (source-transport) was almost useless for prioritization, as it identified all plots to have similar risk. However, the component-flow based index (surface runoff + subsurface runoff + erosion) resulted in sharp prioritization (2.5% high risk, 15% medium risk), with similar risk shares to APLE modeling (i.e. 18% contributing to 50% of P loss). The congruence between component-flow indices and APLE has also been found in earlier studies (Fiorellino et al., 2017; Osmond et al., 2017). The two approaches provide complementary views to the same problem because they include different processes. The P index does not include stratification and dissolved P loss. APLE does not include subsurface P loss and calculates runoff from daily rainfall, which in our case underestimated runoff from prolonged wet periods. Overall, P indices and models are promising for field-level priority setting, but their application in the context of wet soils and widely applied conservation tillage would require further development in the runoff and stratification modules. Finally, access to high-resolution erosion maps can help focus on within-field hotspots. In our dataset, 4-13% of the analyzed field area had severe erosion. This is similar to the 1-6% of land area in other catchments that have been identified as critical source areas for P loss (Thomas et al., 2016), further supporting the idea of targeting mitigation to within-field P loss hotspots.

In conclusion, participatory on-farm experimentation is emerging as a key tool for developing more sustainable agricultural practices. In this Carbon Action experiment, volunteer farmers developed and maintained a carbon farming experiment on their fields for five years, keeping detailed notes of their practices and allowing sampling and on-site measurements. The experiment was tailored to match C sequestration processes; therefore, it was not surprising that it failed to simultaneously mitigate P loss. Achieving simultaneous reductions in P loss and gains in C would require on-farm experiments that focus on key processes of P loss: hotspots, removal of legacy P, and reducing runoff. Without targeting these, carbon farming as such is likely to have only a minimal impact on reducing P losses.

## ACKNOWLEDGEMENTS

This work was funded by the Academy of Finland Strategic Research Council grant MULTA (352437). The authors would like to thank Elisa Vainio for comments on the manuscript structure and the participating farmers for enabling the research on their fields.

